# Phosphate starvation induces replacement of phospholipids with the betaine lipid diacylglycerol-N,N,N-trimethylhomoserine in the human fungal pathogen *Candida albicans*

**DOI:** 10.1101/199133

**Authors:** Surabhi Naik, Rebecca Cahoon, Bridget Tripp, Christian Elowsky, Sophie Alvarez, Kan Liu, Chi Zhang, Wayne Riekhof

## Abstract

We have previously demonstrated that phosphate starvation induces replacement of phosphatidylcholine with the betaine lipid diacylglyceryl-*N*,*N*,*N*-trimethylhomoserine (DGTS) in fungi. In *Neurospora crassa*, the *BTA1* gene encodes the betaine lipid synthase, which is necessary and sufficient for DGTS synthesis. BTA1 expression and DGTS accumulation are part of the fungal phosphorus (P_i_) deprivation (PHO) regulon, mediated by the NUC-1/Pho4p transcription factor. We now demonstrate that the human pathogen *Candida albicans* encodes a *BTA1* ortholog (*Ca*BTA1), which is activated during P_i_ scarcity. The *Ca*BTA1 gene is also induced under certain biofilm-promoting conditions independent of P_i_ starvation. RNA-seq and qRT-PCR showed a significant increase in *Ca*BTA1 expression in response to P_i_ limitation. Thin-layer chromatography and LC-ESI-MS/MS confirmed the replacement of PC with DGTS in wild-type under low P_i_ and showed the absence of DGTS in the *bta1ΔΔ* mutant.

P_i_ limitation in the gut of critically ill patients also triggers the switching of *C. albicans* into an invasive filamentous form. To assess the role of BTA1 and DGTS in the pathogenicity of *C. albicans in vitro*, we compared the growth and morphology of *bta1ΔΔ* and wild type in hyphae-inducing media and observed defects in biofilm formation and invasive growth in the *bta1ΔΔ* mutant. This observation is complemented by RNA-seq data demonstrating that P_*i*_ starvation in planktonic *C. albicans* cells induces the expression of virulence-associated cell surface proteins. Taken together, these results show novel functional interactions between lipid metabolism and remodeling, biofilm formation, and the phosphate starvation response of *C. albicans*.

## Introduction

Microorganisms must rely on their external environment for nutrients. Microbes can experience rapid shifts from nutrient excess to defiency, and this environmental variation leads to the adjustment of regulatory mechanisms for optimal growth and survival (1). Phosphorus, in the form of inorganic phosphate (PO_4_^2−^; P_*i*_) is an essential macronutrient for all organisms, and is necessary for the biosynthesis of lipids, nucleic acids, cofactors, and phosphorylated metabolites (2), and a number of strategies have evolved to cope with limited P_*i*_ availability. We have shown previously (3) that P_*i*_ starvation in fungi (*Neurospora crassa* and *Kluyveromyces lactis*) leads to the replacement of phosphatidylcholine (PtdCho) with the phosphorous-free betaine lipid diacylglyceryl-N,N,N-trimethylhomoserine (DGTS). In ascomycetes, this process is under the control of the NUC-1/Pho4p transcription factor which regulates a suite of genes associated with phosphate scavenging and uptake, typically referred to as the PHO regulon (3). We now demonstrate that the biochemical pathways responsible for the turnover of phospholipids and biosynthesis of DGTS in *Candida albicans* are coordinately regulated as a core component of the PHO regulon of this pathogenic fungus. The betaine lipid synthase of *C. albicans* (C1_11490C; *Ca*BTA1) is part a cluster of coregulated genes under P_*i*_ limitation (Fig 1.)

Romanowski *et al.* (4) demonstrated that under physiological stress, the gut of critically ill patients develops severe P_*i*_ depletion which leads to the switching of *C. albicans* from its commensal yeast form to an invasive filamentous form. This finding prompted us to study the morphological switching and virulence in *C. albicans* in relation to its biochemical ability to synthesize DGTS or to replace PtdCho with DGTS in response to phosphate limitation.

*C. albicans* is an opportunistic fungal pathogen present in the healthy human gut microbiota. Immunocompromised or immunosuppressed patients, particularly AIDS patients, cancer chemotherapy patients, organ transplant recipients, and neonates are more susceptible to *Candida* infections/candidiasis (5). *C. albicans* is referred to as a dimorphic fungus, because it can switch from an avirulent commensal form to a virulent invasive form in response to various environmental cues such as nutrient starvation, pH, temperature, CO_2_, and adherence. *C. albicans* has the ability to transition between yeast, pseudohyphae and a virulent true hyphal form (6–9). A major factor of *C. albicans* virulence is its ability to form biofilms consisting of densely colonized yeast and hyphal cells adhered to a surface (10, 11). Biofilm development consists of an initial adherence phase, followed by filamentous growth and maturation in which an extracellular matrix accumulates, making biofilm-associated *Candida spp.* infections highly resistant to conventional antifungal drugs and the host immune system. Upon maturation, the hyphae can resume growth as non-adherent yeast cells, facilitating further dissemination (10, –14). While many studies, e.g. (8, 15–17), have investigated the regulation of morphological switching in *C. albicans*, little is known about how P_*i*_ limitation promotes hyphal morphogenesis, or whether this stress induces changes in lipid composition, as we have shown for other fungi.

Our current study demonstrates that: i. P_*i*_ limitation induces the degradation of phospholipids and their replacement with the P_*i*_-free lipid DGTS; ii. Transcriptomic changes under P_*i*_ limitation suggest an increased flux of carbon from serine and glycine into the C1-pathway and increased flux into methionine, S-adenosylmethionine, DGTS, and polyamines; iii. The BTA1 gene is expressed in certain biofilm inducing conditions regardless of P_*i*_ nutritional status; and iv. Deletion of the BTA1 gene results in modest defects in biofilm formation under P_*i*_ replete conditions, and severe defects in biofilm formation when P_*i*_ is limiting. Taken together, these results demonstrate novel connections between the P_*i*_ starvation response, hyphal growth, biofilm formation, and amino acid metabolism, and provide new insights regarding biochemical adaptations to P_*i*_ starvation that might be exploited in the development of new antifungal compounds.

## RESULTS

***Pathways associated with phospholipid degradation and DGTS synthesis are induced by P*_*i*_ *starvation in C. albicans.*** The biosynthesis of the betaine lipid diacylglyceryl-*N,N,N*-trimethylhomoserine (DGTS) has previously been studied in various organisms including the purple bacterium *Rhodobacter sphaeroides* (3) and green alga *Chlamydomonas reinhardtii* (18), and it has been observed that this non-phosphorous glycerolipid (DGTS) replaces phosphatidylcholine (PtdCho) during phosphorous starvation. Expression of DGTS synthases in *E. coli* and *S. cerevisiae*, both of which lack the capacity to synthesize this lipid, results in DGTS accumulation. We have previously shown that BTA1 is necessary and sufficient for DGTS synthesis under P_*i*_-limiting conditions in *N. crassa*, and is under the control of the PHO regulatory system transcription factor NUC-1 (3).

We identified a BTA1 homolog in *C. albicans*, and given the genomic context of the BTA1 gene, coupled with RNAseq data for P_*i*_ starved cultures, we postulate a regulatory network of DGTS synthesis. Fig. 1A shows a 35kb region of *C. albicans* chromosome 1, which contains a cluster of genes which we predicted to be coordinately regulated for the synthesis of betaine lipid in response to phosphate limitation. DGTS biosynthesis is regulated as part of the PHO regulon, which controls the expression of BTA1 and is likely to regulate the activity of other members of the cluster, including SAM2, PLB1, GDE1, GPT2, SLC1, PAH1, and PHO84, which were induced as determined by our RNAseq analysis. We propose that a sequential biochemical pathway synthesizes DGTS, using fatty acids and glycerol-3-phosphate released from PtdCho and PtdEtn degradation as substrates. We propose that PtdCho is deacylated by the PLB1 enzyme, which is the predominant phospholipase B isoform of *C. albicans* (19), responsible for the synthesis of fatty acid and water-soluble glycerophosphocholine (GroPCho). The second step is catalyzed by the glycerophosphodiesterase, GDE1, which hydrolyzes GroPCho to choline and glycerol-3-phosphate (Gro-3-P). Glycerol-3-phosphate acyltransferases (*Ca*GPT2 or SCT1) acylate glycerol-3-phosphate at the *sn-1* position yielding lysophosphatidic acid (lyso-PA), which then further gets converted into phosphatidic acid (PtdOH) by acylation in the sn-2 position. This reaction is catalyzed by a 1-acyl-Gro-3-P acyltransferase (AGPAT) or, lysophospholipid acyltransferase. The genes known to encode such an AGAT are SLC1 and *Sc*ALE1/*Ca*LPT1 (lysophospholipid acyltransferase) (20, 21). Dephosphorylation of PtdOH is catalyzed by a phosphatidate phosphatase (*Ca*PAH1) yielding diacylglycerol (DAG). Nakamura *et al.* reported that eukaryotic Pah1 is responsible for membrane lipid remodeling under phosphate starvation (22). It has been reported that under low phosphate conditions, Pho85p-Pho80p phosphorylates Pah1p thereby regulating phosphatidate phosphatase activity (23). Cellular trafficking of Pah1 plays a key role in PtdOH dephosphorylation. Firstly Pah1 is phosphorylated by Pho85-Pho80 complex in the cytosol, resulting in its recruitment into the nuclear/ER membrane, where it gets dephosphorylated by the Nem1p-Spo7p protein phosphatase complex. This dephosphorylation allows the attachment of Pah1 to the membrane where it dephosphorylates PtdOH into DAG (24). We have previously demonstrated the synthesis of DGTS from DAG by BTA1 in bacteria, genetically engineered *S. cerevisiae*, and the fungi *K. lactis* and *N. crassa* (3). *Candida albicans* encodes a BTA1 ortholog (*Ca*BTA1), which is activated during phosphate limitation and is responsible for DTGS synthesis. SAM2, which encodes for S-adenosylmethionine synthetase, synthesizes S-adenosylmethionine (AdoMet) from methionine and ATP, and is highly induced under P_*i*_ limitation. AdoMet acts as the donor of the four-carbon homocysteine/homoserine carbon skeleton to the DAG molecule, and as a methyl donor for N-trimethylation of the diacylglycerylhomoserine intermediate, resulting in the synthesis of DGTS by BTA1. This regulatory network is further validated by analytical detection methods suggesting that switching of PtdCho into DGTS in phosphate scarce condition.

**Fig. 1.**
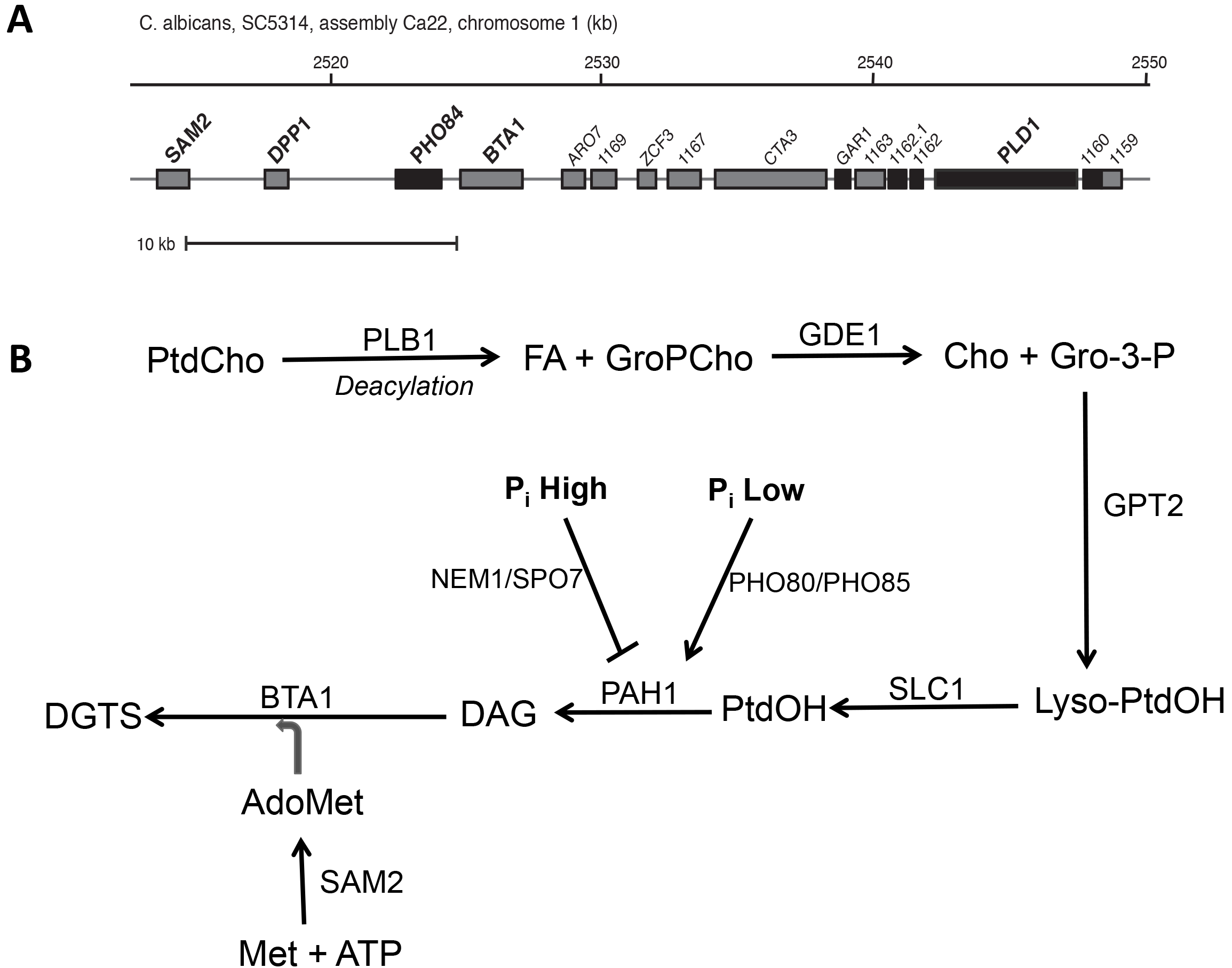
Schemetic of a proposed regulatory pathway of phospholipid turnover and betaine lipid synthesis in *C. albicans* under phosphate limitation. (A) a 35 kb region of *C. albicans* chr. 1, consist of co-regulated genes responsible for betaine lipid synthesis. (B) Pathway showing most likely phospholipid to DGTS pathway in *C. albicans* under P_*i*_ limitation. Abbreviations: PtdCho, phosphatidylcholine; GroPCho, glycerophosphocholine; FA, fatty acid; G-3-P, glycerol-3-phosphate; Lyso-PA, Lyso-phosphatidic acid; PtdOH, phosphatidic acid; DAG, diacylglycerol; DGTS, diacylglyceryl-N,N,N-trimethylhomoserine; AdoMet, S-adenosylmethionine

*Identification of DGTS under P*_*i*_ *starved growth conditions*

To confirm the function of this regulatory pathway and the expression of *Ca*BTA1 under phosphate limitation, we performed lipid analysis by thin layer chromatography (TLC) followed by electrospray ionization tandem mass spectrometry (ESI-MS/MS) analysis of isolated lipid species. Identification and characterization of DGTS in the wild type SN152 and *bta1*ΔΔ mutant (FGSC) was done by ESI-MS/MS. Under P_*i*_ limited conditions, a decrease of PtdCho and a corresponding quantitative increase of DGTS in the wild type was observed. The absence of DGTS in the *bta1*ΔΔ mutant was confirmed by TLC (Fig 2A) and individual bands of DGTS and PtdCho were isolated and further characterized by direct infusion electrospray ionization-MS/MS with neutral loss of 236.1 and precursor ion scan of 184.1 to confirm PtdCho and DGTS molecular species respectively (Fig. 2B).

ESI-/MS/MS scans were quantified and molecular weights of individual molecular species were used to calculate the fatty acid composition of PtdCho and DGTS (Fig 2C), and gas chromatography of fatty acid methyl esters was used to assess the overall cellular fatty acid profile (25)(Fig. 3.) A significant difference in the composition of molecular species of DGTS and PtdCho was detected under phosphate limited conditions. The major molecular species of DGTS were 32:1 (total carbon atoms:number of double bonds), 34:1 and 34:2 along with some of the odd chain fatty acid as shown in Fig. 2C. The most abundant molecular species of PtdCho present were 34:2, 36:3, 36:4, 36:5 and some odd chain fatty acid (35:2, 35:3, 37:2 and 37:3).

The fatty acid profile of DGTS shows that it primarily consisted of 32 and 34 carbon molecular species, whereas PtdCho consists of 34 and 36 carbon unsaturated fatty acid (Fig. 2C). Several studies have been reported the importance of the balance between the saturated and unsaturated fatty acid for maintaining the fluidity, cell integrity, pathogenicity and optimal functioning of the cell. Thus we further examined the fatty acid composition of the WT and *bta1ΔΔ* whole cells lipid extracts under phosphate limited conditions to assess the unsaturation of each fatty acid class in Fig. 3A.

The unsaturation index was calculated by taking the sum of the fatty acyl chain mole percentage, multiplied by the number of double bonds in that species (26). The unsaturation indices of *C. albicans* wild types and *bta1*ΔΔ fatty acid show a significant decrease in unsaturation under phosphate-limited conditions because of lower levels of 18:2 and higher levels of 18:1. This result correlates well with our gene expression data, which indicated increase in OLE1 (stearoyl-CoA desaturase) expression and a significant increase in FAD2 and FAD3 expression in wild type, in response to phosphate starvation. Fig. 3B shows the ratio of 16 and 18-carbon acyl chain, showing the presence of more 18-carbon fatty acid under phosphate starvation. The unsaturation indices of PtdCho were significantly higher than DGTS indicating that DGTS had fewer double bond per acyl chain.

**Fig. 2a.**
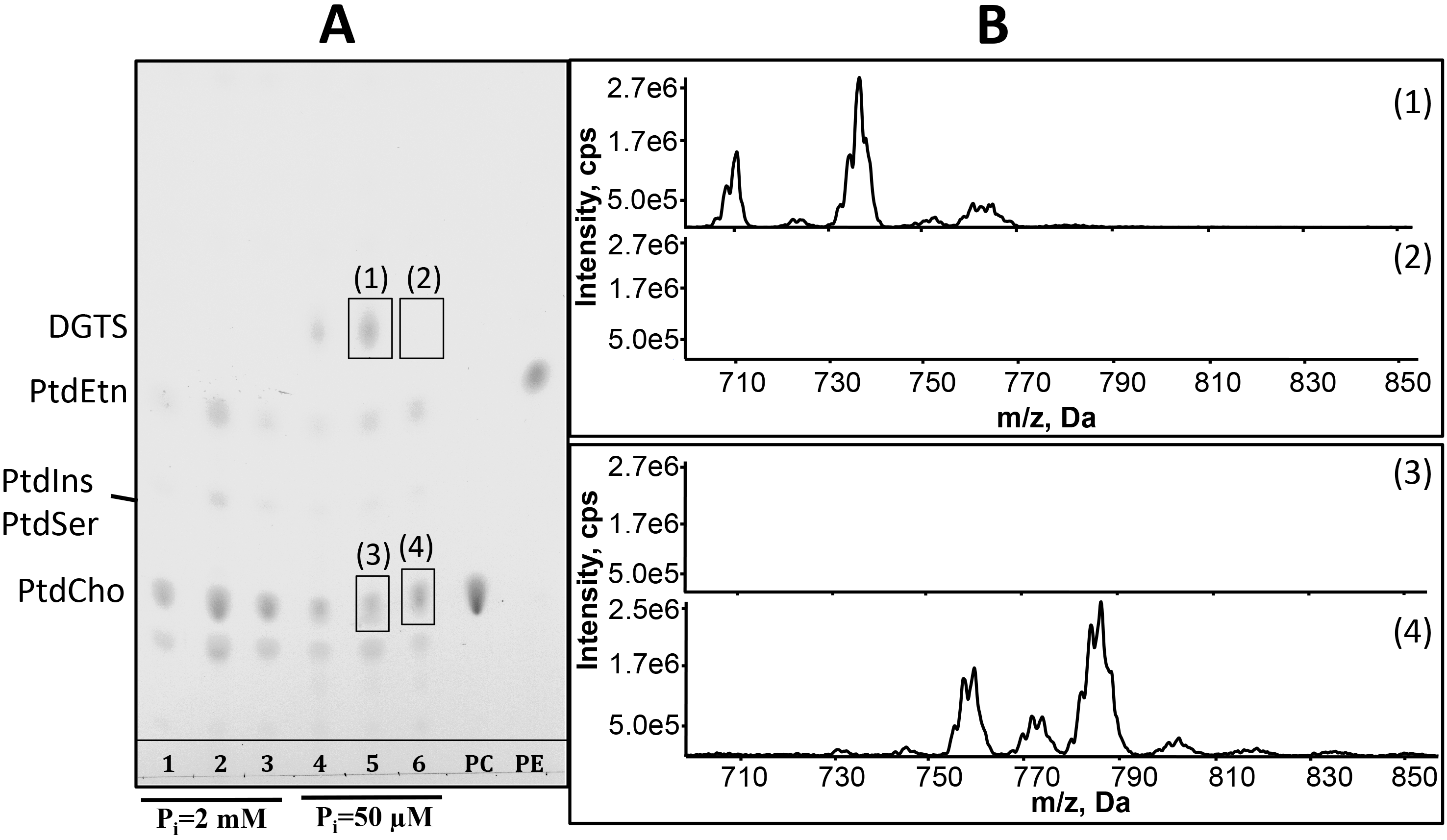
Replacement of PtdCho with DGTS in wild type and *bta1ΔΔ* under phosphate limited (50μM) and replete (2mM) conditions. Identification of DGTS by (A) Thin Layer Chromatography where 1,4 - SC5314; 2,5 - SN152; 3,6 - *bta1ΔΔ*; PC, phosphatidylcholine and PE, phosphatidylethanolamine. (B) Electrospray ionization mass spectrometry (ESI-MS/MS). (1, 3) Chromatogram showing the presence of DGTS and absence of PtdCho in WT respectively and (2, 4) Chromatogram showing the presence of PtdCho and absence of DGTS in WT under P_i_ limited condition in the bta1ΔΔ mutant strain. (C) Molecular species quanitification of DGTS and PtdCho analyzed by ESI-MS/MS.

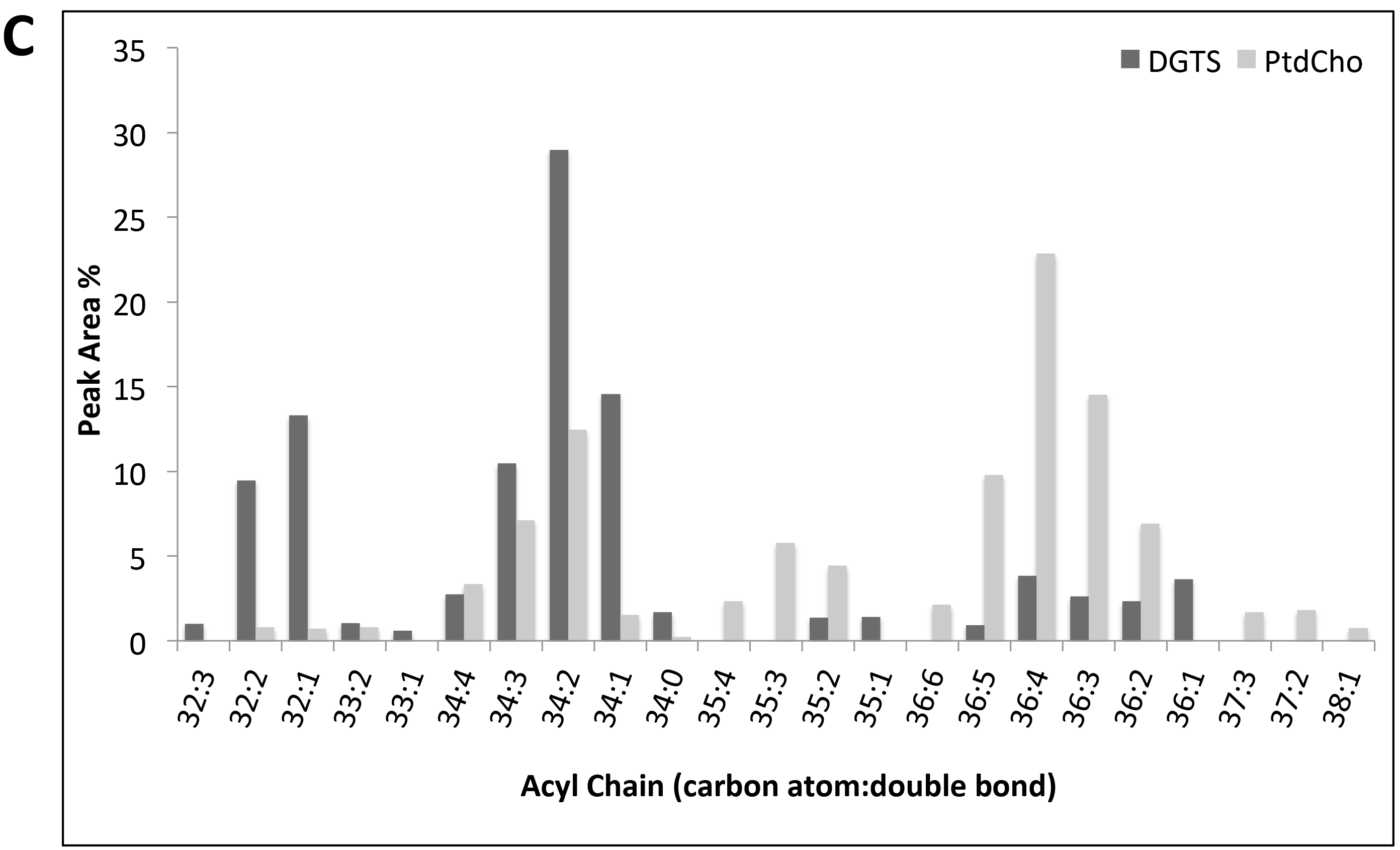

**Fig. 3.**
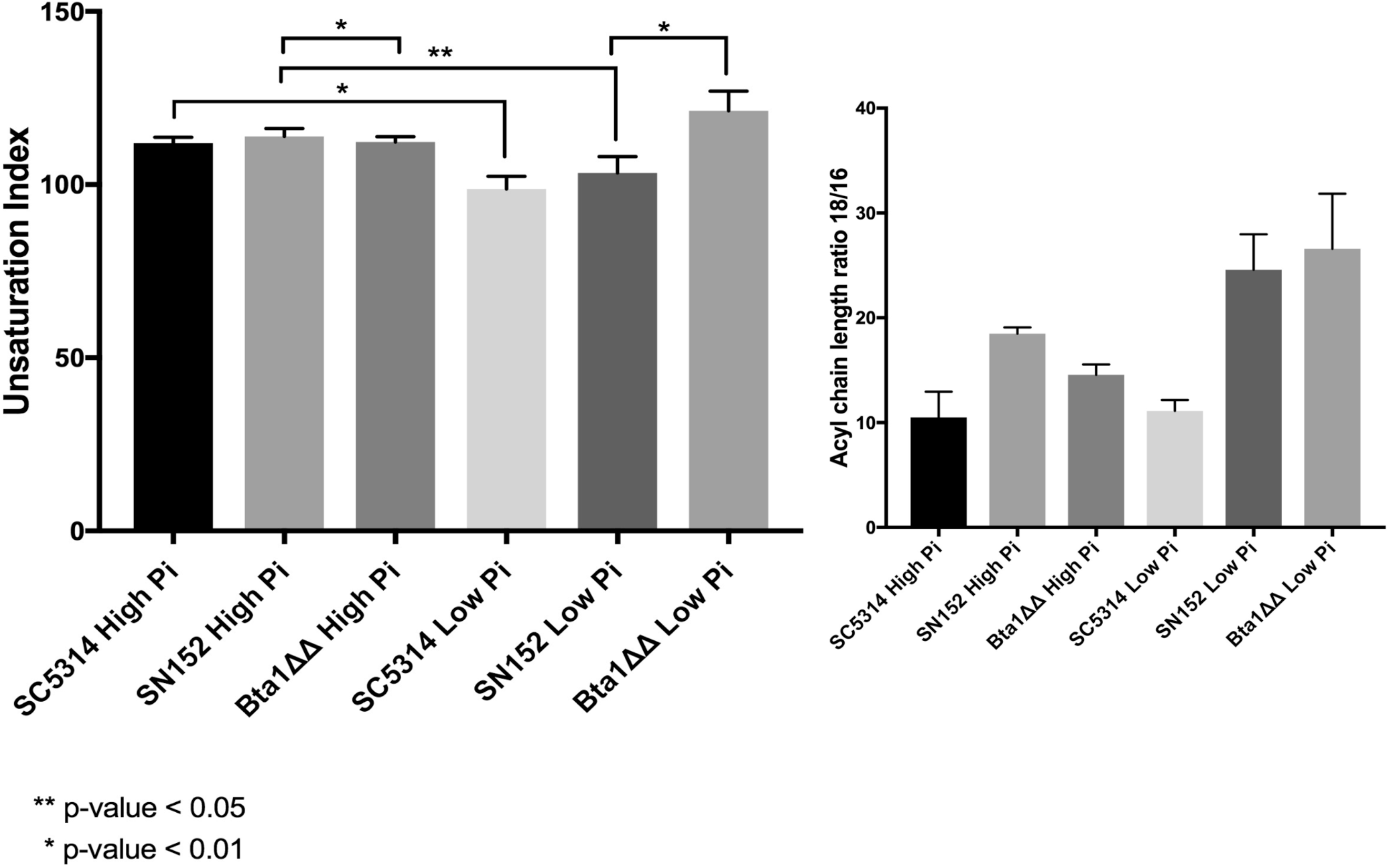
Comparative fatty acid composition in wild type (SC5314 and SN152) and *bta1ΔΔ* under phosphate limited and replete condition. (A) Unsaturation Index of each fatty acid class, (B) Ratio of 16-carbon acyl chain and 18-carbon acyl chain fatty acid.

We then verified that the expression of BTA1 which is induced in response to P_i_ limitation and is under the control of PHO regulon, by performing quantitative real-time PCR (qPCR) analysis as shown in figure 4. We observed a significant increase in BTA1 and PHO84 expression under P_i_ limited conditions. In addition, we also checked the changes in the expression level of other genes in the pathway; PLD1, DPP1, and SAM2 in wild type under phosphate limited and replete conditions and no significant fold change were observed.

**Fig. 4.**
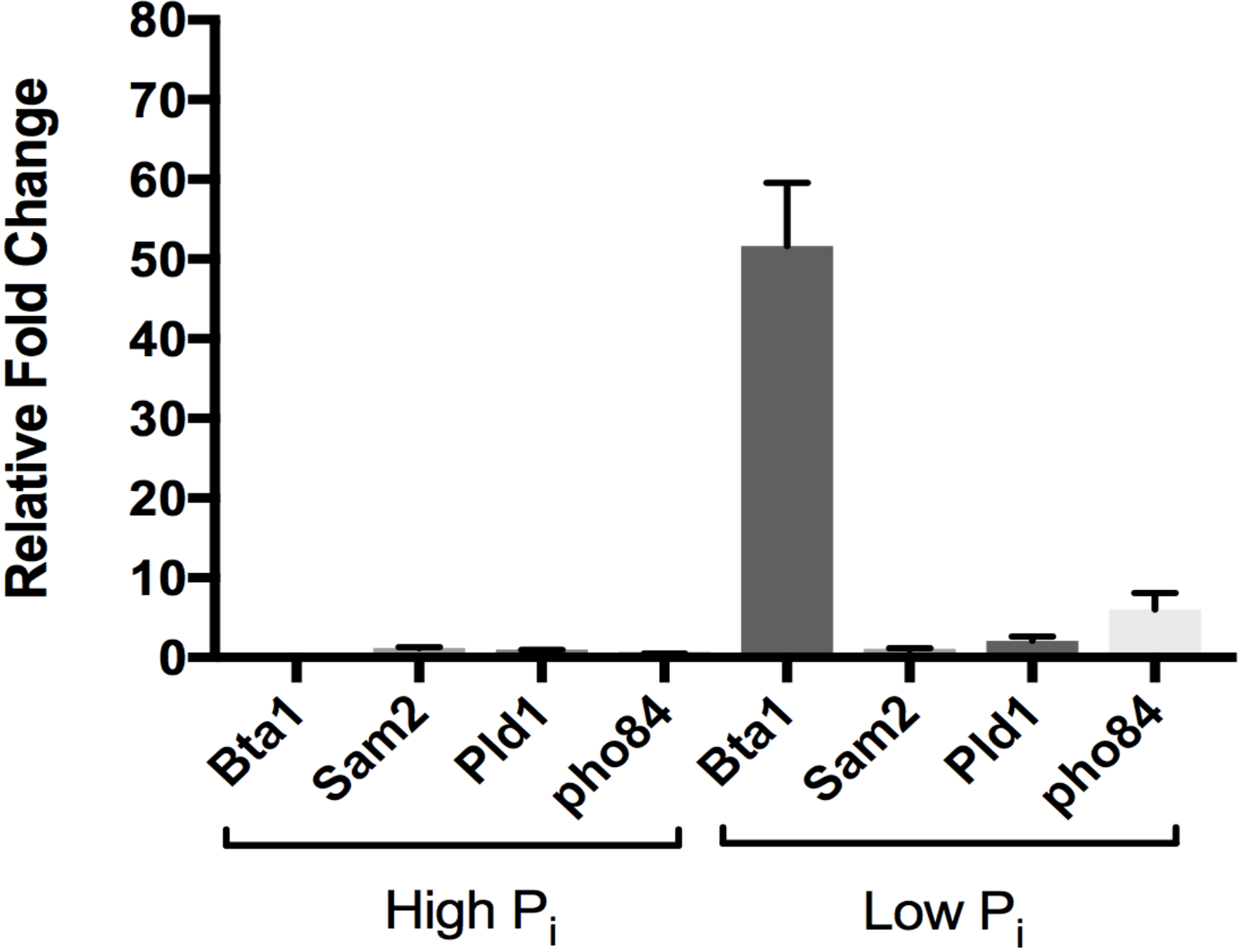
Gene expression of BTA1, SAM2, DPP1, PLD1 and PHO84 in wild type under P_i_ limited (50 μM) and replete (2 mM) conditions grown in synthetic complete media, by quantitative real time PCR (qRT-PCR). Data represent relative expression levels as normalized to *Ca*ACT1 as an internal control.

### Role of *bta1* in colony morphology and hyphae formation

It has been shown that *Candida* hyphae are capable of invading both agar and epithelial cells *in vivo*. To determine the role of BTA1 in hyphae formation, the wild type and *bta1*ΔΔ mutant were grown on hyphal-inducing media (solid agar plates as well as liquid media). Strains were cultured in YPD media overnight at 30°C, then washed twice with the media and spread with 1:10 dilution on spider, serum (10%), lees and YPD agar plates. Culture plates were incubated for 48-72 hours at 37°C and photographed (Fig. 5). We observed that there was no significant difference in the colony morphology of wild type and *bta1*ΔΔ mutant when grown in serum agar media, whereas the wild type forms more clear hyphae than *bta1*ΔΔ mutant on spider and lee’s media. When grown on solid spider media, wild type colonies develop an expanded central wrinkle region, which is a mixture of hyphae and pseudohyphae, and an invasive peripheral filament region. The *bta1*ΔΔ mutant has no difference in invasive growth, but lacks expanded aerial hyphae.

**Fig. 5.**
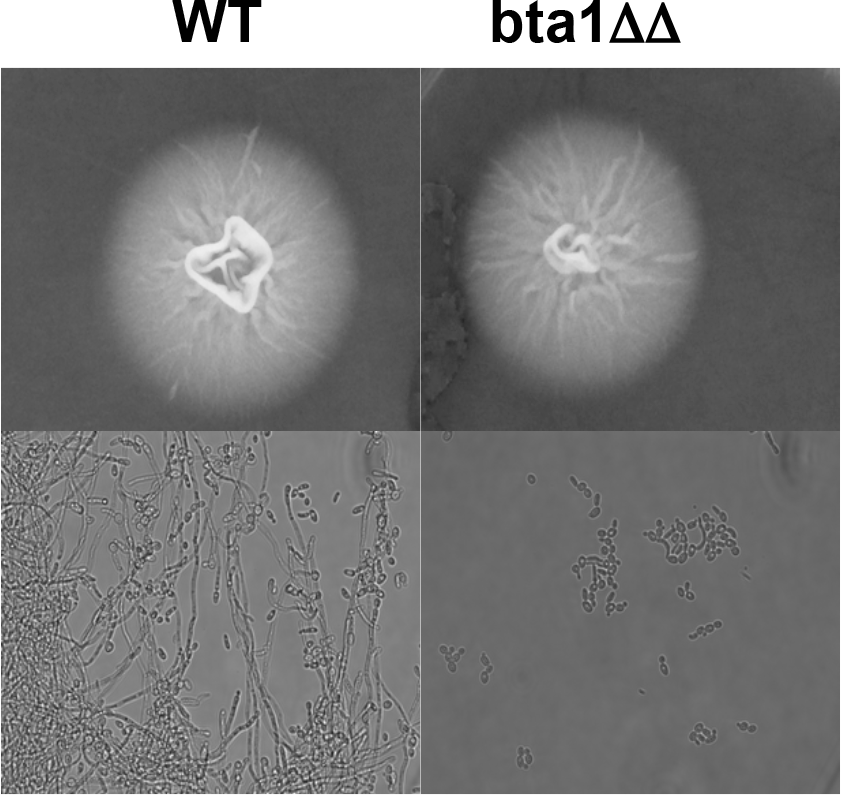
Colony morphology of wild type and *bta1*ΔΔ grown on (A) Spider agar plates (B) Spider liquid media.

To further investigate the role of BTA1 in filamentation we grew the strains in spider liquid media and incubated for 48 hours at 37ΰC Wild type showed distinctive long, invasive filaments whereas *bta1*ΔΔ mutant showed the presence of pseudohyphae. Taken together, these results suggest that BTA1 might be required for optimal invasive hyphae formation.

### Role of BTA1 in Biofilm formation

Fungal hyphae are a key feature in biofilm development but not necessarily essential for biofilm establishment or maintenance. Rather it plays a role in strengthening the biofilm structure, which leads to resistance to antifungal drugs. Development of *C. albicans* biofilms consist of four steps: first is adherence of yeast cells to the solid surface (adherence step), second is initiation step in which adhered cells starts to grow and form elongated hyphae or pseudohyphae. The third is maturation step in which long filaments begins to secrete extracellular matrix (ECM) leading to increased drug resistance. The final step is dispersal step in which oval yeast cells are released into the external environment, leads to disseminated infection (13).

To determine the morphology and architecture of biofilm, the wild type, and *bta1*ΔΔ mutant were pre-grown in YPD media overnight at 37°C. Small square silicon pieces were cut from silicon sheet and washed with sterile water followed by autoclaving. Sterile silicon pieces were then incubated overnight with fetal bovine serum and then washed with media immediately before inoculation. For the initial adherence step, inoculation was done with initial OD 0.5 in 2 ml RPMI-1640 media containing silicon pieces in 12 well plates, followed by incubation at 37°C for 90 min with gentle shaking (150rpm). Each silicon piece was washed gently with media, and fresh RPMI-1640 media was added and incubated at 37°C for 60h and 150 rpm for further elongation and maturation steps.

Biofilms were visualized after 24h and 48h time interval to study biofilm morphology of initial biofilm and mature biofilm. Biofilms were stained with calcoflour white stain for visualization in confocal laser scanning microscopy (CLSM). Depth views were conducted with artificial color gradient indicating cells closest to silicon piece in blue and furthest from the substrate in red. We compared the ability to form a stable biofilm in the wild type and *bta1*ΔΔ mutant in-vitro, under phosphate limited and replete conditions. It was observed that the biofilm structure varies significantly under P_i_ limitation. To investigate the role of BTA1 in the structure of biofilm, we grew the cultures in different media conditions (RPMI 1640, spider, lee and serum media). It was observed from the microscopic view of the biofilms that the wild type forms very intact biofilm whereas *bta1*ΔΔ mutant form an attenuated and impaired architecture of biofilm in RPMI media (Fig. 6B). The depth of the biofilm was measured as 26 micron for wild type and 85 micron for the *bta1ΔΔ* mutant. We found that the *bta1ΔΔ* mutant form mixed biofilm, which consists of planktonic yeast cell and hyphal/pseudohyphal forms, in lee’s, spider and serum media.

It has been previously elucidated that chitin-glucan linkage is essential for the control of morphogenesis in *candida albicans* (27). Hence to investigate the role of BTA1 in chitin synthesis of *candida albicans*, we measured the chitin accumulation in wild type and *bta1ΔΔ* mutant, by calcoflour white staining. Calcoflour white is used to stain and visualize chitin in the yeast cell wall. A chitin ring is formed at bud emergence at the neck between mother and daughter cell, which is visualized by calcoflour white fluorescence assay. We observed a significant difference in the calcoflour white staining in wild type and *bta1ΔΔ* mutant. Deletion of *bta1* results in the decrease of chitin accumulation at the septum of *bta1*ΔΔ mutant as shown in fig. 6A.

**Fig. 6.**
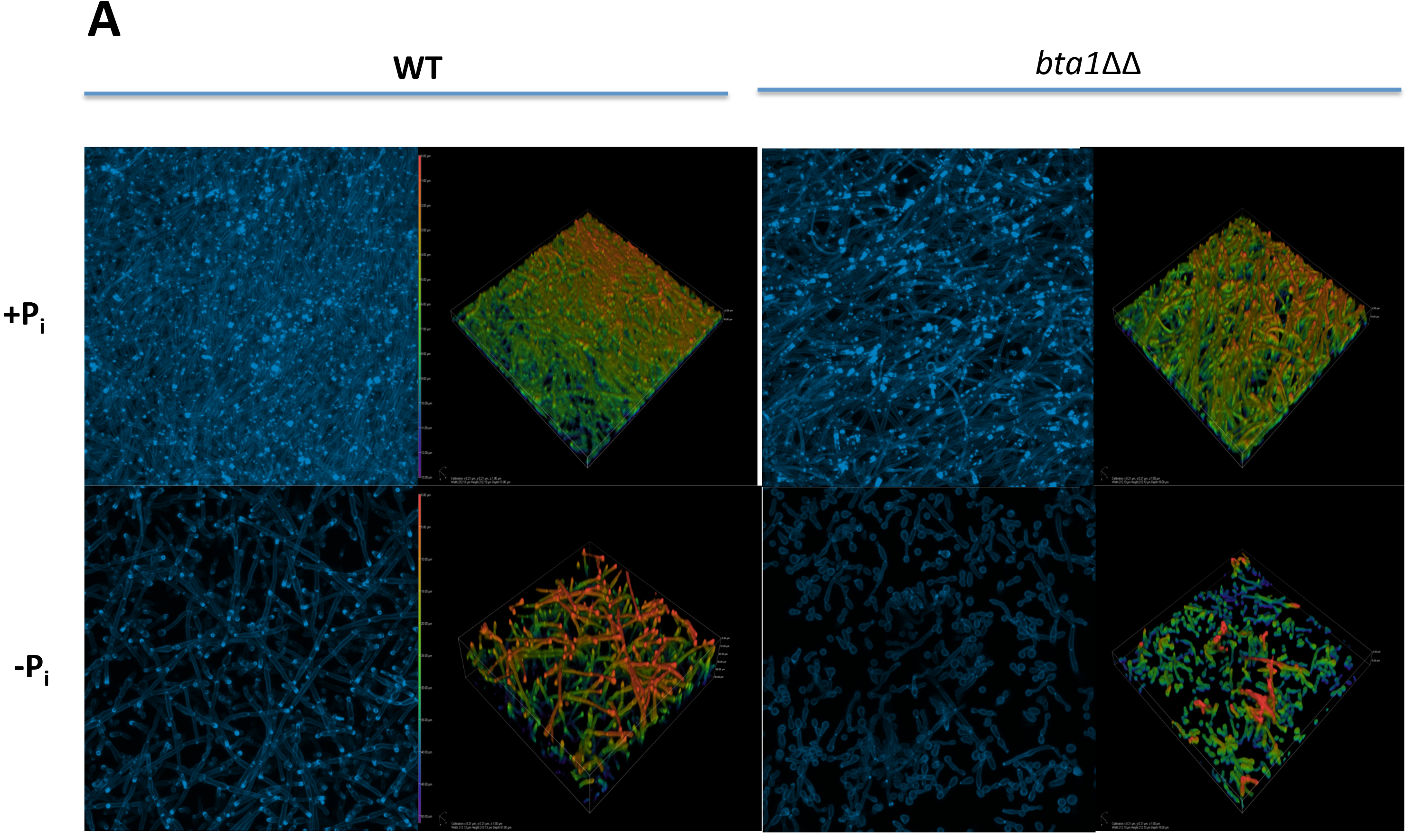
Confocal laser scanning microscopy (CLSM) images show altered filamentation and biofilm architecture of *bta1ΔΔ C. albicans in vitro*. (A) 3D images of WT and *bta1*ΔΔ biofilms grown under P_*i*_ replete and limited conditions, stained with calcoflour white, and imaged by CLSM. (B) Depth view of WT and *bta1*ΔΔ biofilm grown under different media (RPMI-1640, Lee’s media, spider and serum media).

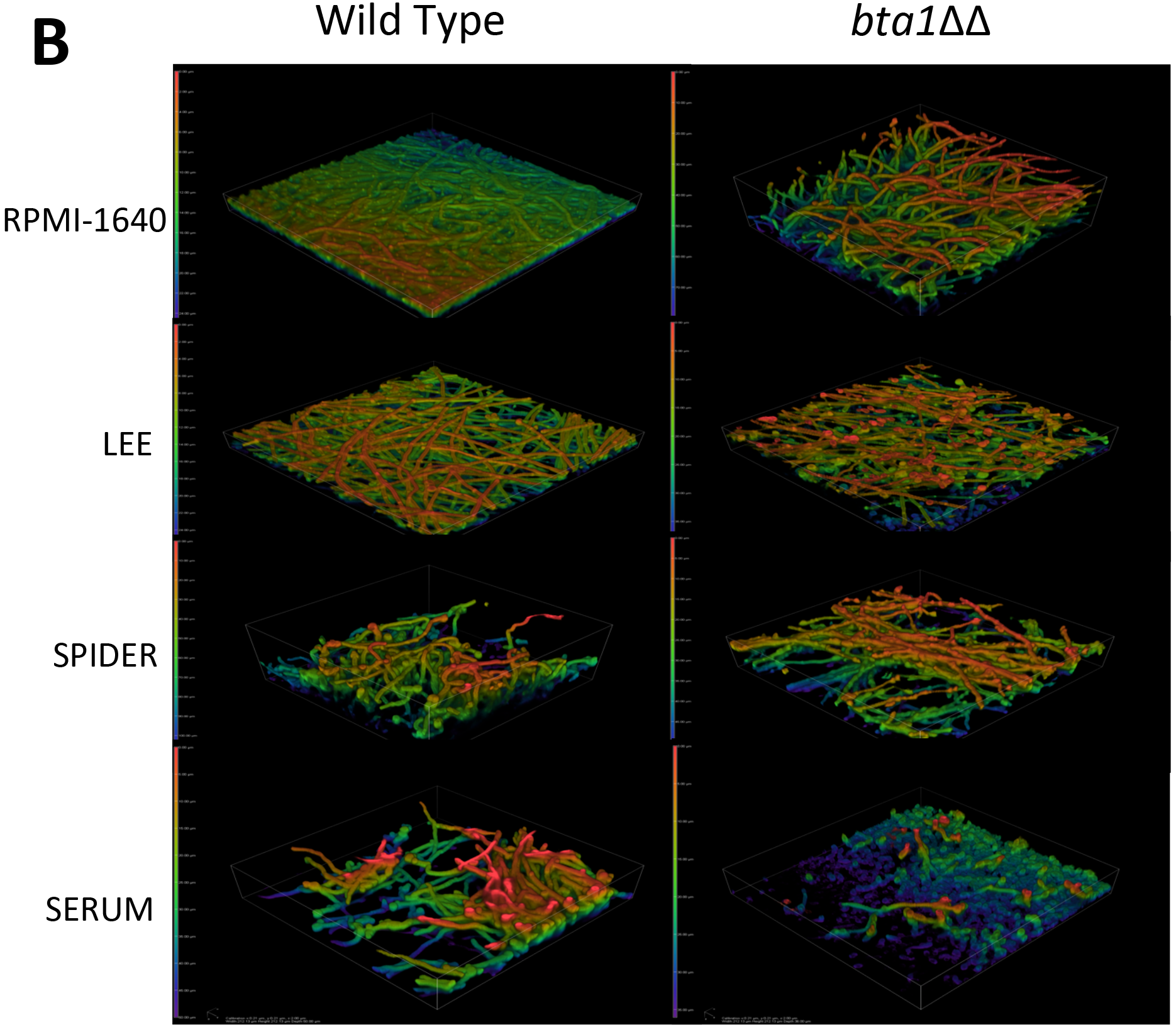

### RNA-Sequencing Analysis and validation by qRT-PCR

In addition to microscopy, phenotypic and biochemical analysis, we performed RNA-sequencing based transcriptional profiling to determine the role of BTA1 and DGTS in maintaining cellular lipid homeostasis, biofilm formation, hyphal formation, iron and copper homeostasis and pathogenicity, in response to P_*i*_ starvation in *C. albicans*. Wild type SN-152 and *bta1*ΔΔ mutant strains were used for differential gene expression studies. Comparable transcriptional regulation was done under phosphate deplete and replete conditions. We observed that BTA1 and PHO84 were highly induced in wild type under P_i_ limited condition, which supports our hypothesis that the BTA1 is expressed under phosphate limited conditions and is a tightly regulated component of the PHO regulon in *C. albicans*, as has been previously reported in *N. crassa* and *K. lactis* (3).

Fig. 7A shows the number of genes conserved, we compared the down-regulated gene expression of WT and *bta1*ΔΔ under phosphate replete condition, it was observed that 69 genes were unique to the wild type and 30 genes were unique to *bta1*ΔΔ sharing 78 common genes. Whereas when upregulated genes were compared 57 genes were unique to the WT and 40 genes were unique to *bta1*ΔΔ sharing 173 genes.

**Fig. 7.**
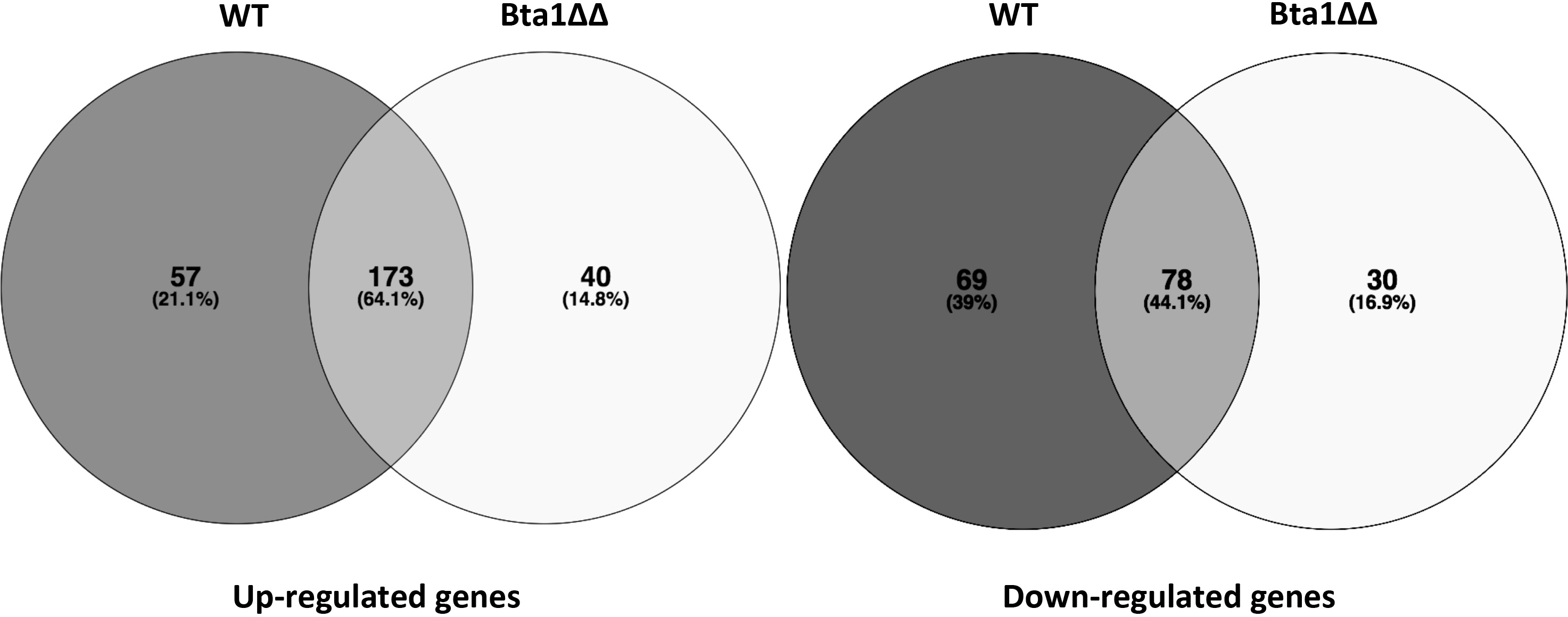
Venn diagram depicting the number of significant (overlapping) differentially expressed genes in wild type and *bta1ΔΔ*. Differential expressed genes were selected on the basis of negative binomial distribution of the count reads using EdgeR, FDR adjusted p-valu<0.05.

An examination of the genes involved in phosphate regulation and storage revealed that high-affinity phosphate transporters PHO84 and PHO89, acid phosphatase genes PHO100, PHO112, and PHO113 were significantly upregulated and low-affinity phosphate transporters PHO87 was significantly downregulated. In addition, we also observed induction of GIT1 which encodes for a permease responsible for utilizing exogenous glycerophosphoinositol and glycerophosphocholine as a P_*i*_ sources (19).

P_*i*_ limitation led to the differential regulation of 377 genes in wild type, and disruption of BTA1 led to differential expression of 321 genes with at least 2-fold change. Of these genes, 230 were upregulated, and 147 were down-regulated in WT, in response to phosphate limitation. To analyze the differentially expressed genes at the functional level, we performed Gene Ontology (GO) and KEGG pathway enrichment analyses using DAVID (Database for Annotation, Visualization and Integrated Discovery; https://david.ncifcrf.gov) to obtain the enriched biological processes (BPs) and pathways using p-value <= 0.05.

Gene Ontology analysis revealed the number of genes involved in biological process and molecular function. The significantly enriched genes from our upregulated gene list were involved in cellular response to drugs, glycine catabolism, ribosome biogenesis, and rRNA maturation and processing. A major role of nutrient signaling is the management of ribosome biogenesis and the translational apparatus in response to nutrient starvation. Biological processes involved in cell adhesion, cellular response to starvation, filamentous growth, pathogenesis, iron and copper transport were significantly enriched in the down-regulated gene list. The down-regulated genes involved in filamentous growth or hyphal morphology were ECE1, ALS1, ECM22, HWP1, QDR1 and SAP2. This result is consistent with the phenotype we have observed for wild type and *bta1ΔΔ* mutant, that *bta1ΔΔ* lacks filamentation and hyphal formation. We also observed genes involved in biofilm formation, and cell adhesion, which were significantly down-regulated.

#### Amino Acid Profiling

Pathway analysis using the KEGG pathway database showed that biosynthesis of free amino acid, glycine, serine, threonine and biosynthesis of secondary metabolites were down-regulated whereas the alanine, aspartate, glutamate, cysteine, methionine and purine metabolic pathways were significantly upregulated in response to phosphate limitation. It was validated by measuring amino acid levels in wild type under phosphate limitation using ultra-performance liquid chromatography (UPLC).

Our amino acid profiling data (Fig. 8) showed a significant decrease in the total free amino acid levels in wild type under P_*i*_ limited conditions. We observed a significant change in the relative concentrations of free amino acid levels in wild type SN152 in response to phosphate limitation. Some amino acid concentration has been decreased specifically e.g. alanine, aspartic acid, lysine, ornithine, serine and increase in glutamine levels were observed in wild type under phosphate limitation.

**Fig. 8.**
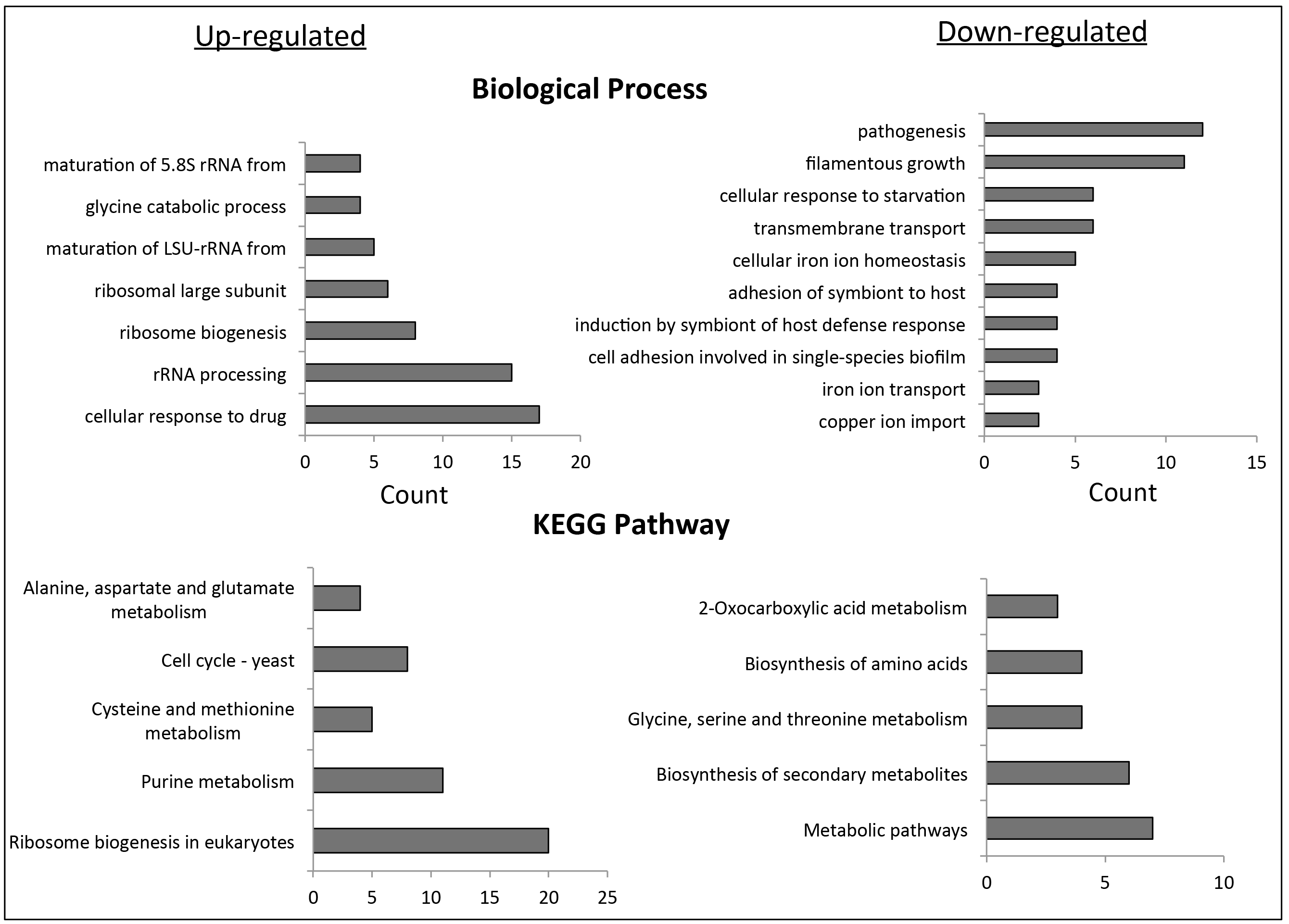
Histograms show significantly enriched Gene term distribution and KEGG analyses with differentially expressed genes in wild type under high and low phosphate condition, to get to obtain the enriched biological processes (BPs) and pathways using p-value <= 0.05. Counts indicate the number of DEGs per category. GO term enrichment and KEGG analyses were performed with DAVID bioinformatics tool. DEG analysis was done by (A) EdgeR software package (B) TopHat and Cufflinks open-source software tools.

In support of the above data, Gene Ontology and KEGG pathway analysis revealed amino acid and ammonium permeases (GAP1 and MEP2) responsible for amino acid and ammonium transport from the plasma membrane were down-regulated in wild type under phosphate-limited condition. It has previously observed that in the presence of the preferred nitrogen source (e.g. glutamine or glutamate or ammonium), Gap1 is repressed transcriptionally and post-transcriptionally (28). GST1, which encodes for glutathione S-transferase, involved in oxidative stress response and cellular detoxification is downregulated. SAP2, a major secreted aspartyl protease considered to show a fundamental role in *Candida* virulence by providing amino acids to the pathogen and degrading host’s proteins was also significantly repressed.
AAT22 encodes for the aspartate aminotransferase, which plays a role in the metabolism of nitrogen and amino acids by catalyzing the reversible transfer of the amino group from L-aspartate to 2-oxoglutarate to form oxaloacetate and L-glutamate (29), is significantly upregulated in wild type under phosphate limitation. JEN1, which is a lactate transporter and also facilitates selenite accumulation inside cells (30), is upregulated in response to phosphate limitation.

*Cellular response to metal ions.* Metal ion such as iron, copper, and zinc are essential nutrients because of the crucial role they play in fundamental cellular processes and also in regulating the pathogenicity of *Candida*. It has been demonstrated that the negative charge in phosphate can bind to the metal cations thereby maintaining phosphate homeostasis in the cell (31). Therefore transcript levels were investigated to understand the phosphate-metal interaction in wild type in under phosphate limitation.

Our RNA-Seq data showed that genes involved in iron, copper, and zinc transport were significantly down-regulated in wild type under low phosphate. Genes belonging to members of the CFL(CFL2, CFL4, FRP1 and FRP2) ferric reductase, the FET (FET34) multicopper ferroxidase, the FTR (FTR2) plasma membrane iron permease family, the iron/zinc-iron transporter (FTH1, RBT5 and ZRT1), as well as copper transporters (AMO2) are down-regulated in wild type under phosphate limited condition. One of the SFU1 (Suppressor of Ferric Uptake) target genes RBT5, which also functionally coordinates with *Ca*TUP1 was down regulated, whereas CCC1, a manganese transporter, was upregulated was in response to phosphate limitation.

## Discussion

Our data suggest some new insights of survival strategies of *C. albicans* under P_i_ limited growth conditions. Metabolic flexibility and adaptability govern the efficient consumption of alternative nutrients in dynamic environment and infection in different host niches. We highlight the membrane lipid remodeling in *candida albicans*, in association with its morphology switching and virulence in various host niches well as signaling pathways. Here we demonstrate that *bta1* gene is solely responsible for the synthesis of betaine lipid in response to phosphate. We show a regulatory pathway of betaine lipid synthesis and the expression of betaine synthase is under the control of fungal PHO regulon.

Our data indicate that the composition of molecular species of DGTS and PtdCho differ in their fatty acid composition and that DGTS undergo a decrease in unsaturation and acyl carbon chain in response to phosphate limitation. It also reveals that DGTS contains fewer double bonds resulting in low cell membrane fluidity that means the cell is less permeable to various ions making it less costly to maintain homeostasis. The maintenance cost can be argued to be one of the key causes of how cell acclimatize in response to stress by increasing the biosynthesis of short-chain less unsaturated fatty acid by decreasing desaturase/elongase activity. It has been reported in many organisms that high unsaturated fatty acids exhibit high membrane fluidity or mobility in response to increasing cold stress (32).

This result indicates that DGTS might be a poor substrate for *ole2*, *fad2* and *fad3*, then PtdCho. Under P_i_ scarce condition i.e. in the absence of PtdCho, *ole2*, *fad2* and *fad3* are highly upregulated, assuming to act on membrane lipid DGTS but because of low substrate specificity with DGTS it results in the loss of 18:2 and 18:3. The unsaturation indices of PtdCho were significantly higher than DGTS indicating that DGTS had less double bond per acyl chain.

In this report, we show that *Ca*Bta1 plays a role in altered morphology as well as its inability to form hyphae, which are reflected in our pathogenicity studies showing inability to produce stable biofilm. Thus the role that *bta1* plays in *C. albicans* is not only in maintaining membrane homeostasis and membrane remodeling for cell survival, but also in the invasive growth response to phosphate starvation. So far the only role known of *bta1* gene is in lipid membrane remodeling under phosphate limitation in other filamentous fungi or bacteria (3, 33), but here we demonstrated its additional role in virulence and pathogenicity in *candida albicans*. It has been reported that adhesion to the substrate is an important early step in biofilm formation. Many studies illustrate that hyphal growth is an important feature and adhesion is crucial during the infection process. Here we show the role of *bta1* in biofilm formation thereby compromising its ability to form a biofilm. Increased levels of cell wall chitin have been known to exhibit reduced sensitivity to echinocandins like drugs (34). Here we exhibit the role of *bta1* in maintaining cell integrity and cellular defense by decreasing the chitin levels. Taken all together, defects in morphology, cell wall, biofilm structure and adherence in *bta1*ΔΔ demonstrate that *bta1* gene has a profound effect on *Candida* defense mechanism and survival.

Our amino acid analysis showed decrease in free amino acid pool in wild type under phosphate deplete condition, consistent with our transcriptome data, which demonstrate the down regulation of amino acid permeases, known to be involved in amino acid transport and metabolism. Our transcriptome data also show the evidence of the crosstalk between transcriptional regulation of phosphate and nitrogen stress response. When P_i_ is limited, the cell drops down its metabolic activity to conserve energy by using critical resources. Under To balance the nitrogen and P_i_ pool inside the cell, it stores glutamate as a nitrogen source as both (N and P_i_) are required for nucleic acid synthesis and imbalance in the ratio could be inhibitory. Our data gives us an idea of how glutamine/glutamate is stored as a nitrogen source (by low expression of ammonium permeases and upregulation of aspartate aminotransferase) under P_i_ limitation.

The defects in phosphate homeostasis have previously been studied in *S. cerevisiae*, pathogenic fungus *C. neoformans* and *C. albicans* (31, 35, 36). It demonstrate that genes involved in iron, zinc and copper acquisition were down regulated but a subsequently increase in the intracellular iron levels in pho4Δ mutant when compared to the wild type, resulting in impaired resistance to metal ions. Our transcriptome data also highlights the connections between phosphate metabolism, metal homeostasis, and superoxide stress resistance for the maintenance of charge stasis in the cell.

Taken together our data demonstrates the biochemical pathway of switching of PtdCho into DGTS, and its transcriptional regulation in response to P_i_ scarce condition. Expression of *bta1* under phosphate starvation can be argued as a preferable low energy demand, cell survival mechanism in response to stress. We conclude that *bta1* gene is not only required for maintaining cellular membrane lipid homeostasis but also for cell pathogenicity under limited nutrient availability. Deletion of *bta1* results in attenuated virulence. We show that *bta1* gene directly or indirectly contributes to the morphology of *candida albicans* and its role in pathogenesis. It elucidates how adaptation and virulence are interconnected in *C. albicans*, but there is still much to be discovered about the mechanisms underlying the adaptability of *C. albicans* within the various host niches.

### Experimental Procedures

#### Strains and growth conditions

*Candida albicans* wild type (SC5314 and SN152) and *bta1*ΔΔ mutant strains were obtained from the Fungal Genetic Stock Center. For phosphate starvation experiments, cells were grown aerobically at 30°C to A_600_ 0.8-1.0 in synthetic complete media (0.67% w/v yeast nitrogen base, 2% w/v glucose, complete amino acid mixture) containing high (2 mM) or low (50 μM) phosphate concentrations. For pre-inoculum, cells were grown aerobically overnight at 30°C in YPD media containing yeast extract 10 g/L, peptone 20 g/L, dextrose 20 g/L, pH-6.5. Cells were harvested by centrifugation and washed twice before inoculation.

#### Lipid Extraction and Analysis

Total lipid was extracted using Bligh-Dyer method (ref.) using chloroform/methanol (2:1, v/v). Briefly, the culture was grown to log phase, harvested and lysed using acid washed glass beads (Sigma). Individual phospholipids were separated by thin layer chromatography (TLC) on silica 60 plates (Sigma-Aldrich) using a chloroform/acetone/methanol/acetic acid/water (50:20:10:10:5 v/v) solvent system. Bands were visualized by brief exposure of the plate to iodine vapor. For identification of individual lipids by ESI-MS/MS, bands were separated by TLC, the silica was scraped into a glass tube, and the purified compound extracted in chloroform/methanol (2:1), and then dried under N_2_ gas.

#### Electrospray ionization-tandem mass spectroscopy (ESI-MS/MS)

Extracted lipids were then analyzed for the phospholipid identification and confirmation using direct infusion ESI-MS/MS approach. Briefly, the extracted lipids were resuspended in 500 μL of chloroform:methanol (2:1) and spiked with 15:0-PC as internal standard. Followed by 5000 times dilution in water/isopropyl alcohol/methanol (55:35:10 v/v/v) containing ammonium formate (25 mM) and 0.4% formic acid, and directly infused into the mass spectrometer at a rate of 10 μL/min. Instrument settings were as follows: source temperature, 300°C; ESI needle voltage, 5.5 kV; declustering potential, 90; entrance potential, 10; curtain gas, 10; gas 1, 50 arbitrary units; gas 2, 40 arbitrary units and Nitrogen gas was used as a collision gas (37).

Lipids were analyzed with a triple quadrupole/linear ion trap mass spectrometer (Sciex QTRAP 4000) in positive ion mode., for precursor scans of 184.1 and 236.1 for PC and DGTS respectively. Phosphatidylcholine was identified with precursor ion scan of m/z 184.1 and DGTS was detected in positive ion mode as neutral loss of 236.1. Scan was taken over the mass range of 200-1000 *m/z* with a cycle time of 2 sec. DGTS showed two peaks at m/z 736.8 and 710.8 with major molecular species of 32:2 and 34:2, in terms of total carbon atoms: double bonds present. In the chromatogram, PtdCho showed four predominant peaks at *m/z* 784.6, 782.6, 758.7 and 757.7 with major molecular species of 36:5, 36:3, 32:3 and 32:1 respectively.

#### RNA isolation and gene expression by qRT-PCR

Cultures were grown to A_600_ 0.8-1.0 on SC-minimal media (described previously) with high (2 mM) and low (50 µM) phosphate concentrations. Total RNA was extracted using OMEGA E.Z.N.A Yeast RNA Kit according to manufacturer’s instructions. RNA purification was done using GeneJET RNA Cleanup and Concentration Micro Kit (Thermo Scientific). cDNA was synthesized using RevertAid First Strand cDNA Synthesis Kit (Thermo Scientific). qRT-PCR was done on Eppendorf Realplex^2^ Mastercycler using SYBR GreenER qPCR Supermix (Invitrogen).

Primer3 software was used to design forward and reverse qRT-PCR primers for *Ca*BTA1, *Ca*SAM2, *Ca*DPP1, *Ca*PHO4 and *Ca*PLD1 (Table I). Gene expression was analyzed using ΔΔC_T_ and normalized to *Ca*ACT1 used as endogenous control. Primers were checked for specificity by melting curve analysis and visualizing in gel electrophoresis. Controls were used lacking the template and reverse transcriptase. Primers for the particular genes did not show any amplification when their respective deletion mutants were used. Wild type low phosphate expression was normalized to their respective internal control (HKG) and then used to compare high phosphate expression. All the experiments were performed in triplicate.

**Table I.**
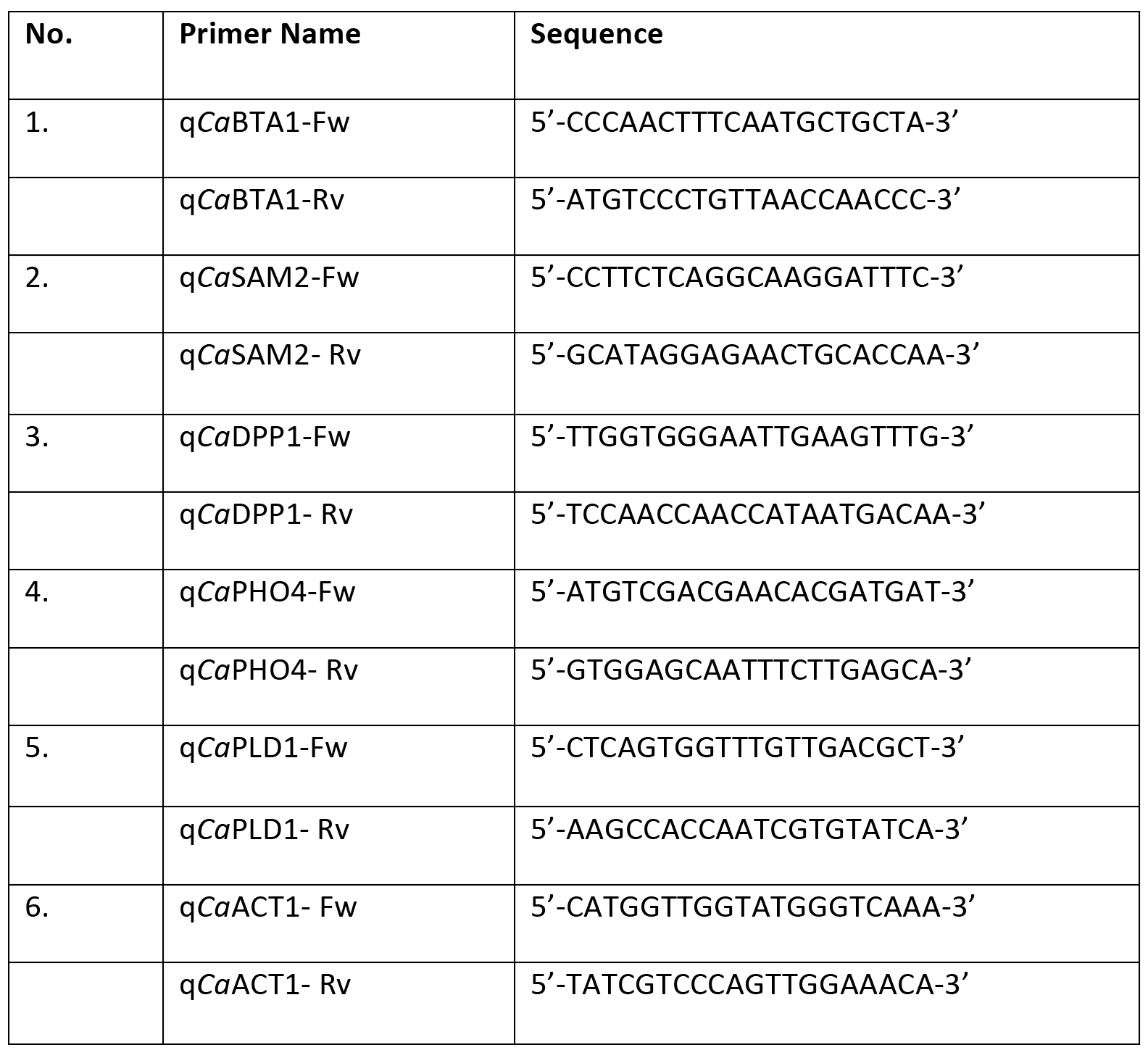
Primers used in this study.

#### RNA-Sequencing Analysis

Briefly, wild type SN-152 and *bta1*ΔΔ mutant were grown to log phase under P_i_ replete and limited conditions Cells were then collected and washed with UltraPure^TM^ DNase/RNase-free distilled water. Total RNA was extracted using OMEGA E.Z.N.A Yeast RNA Kit according to manufacturer’s instructions. RNA purification was done using GeneJET RNA Cleanup and Concentration Micro Kit (Thermo Scientific). RNA samples were sent to Next Gen Sequencing facility, GENEWIZ (www.genewiz.com). Sequencing was performed on the Illumina HiSeq2500 platform in a 1x50bp single-read configuration in Rapid Run mode, with a total of at least 120 million reads per lane or ^~^15M reads per sample. Reads were aligned to the Candida genome (version 22) with Tophat2 (38) and used to calculate differentially expressed genes with the EdgeR software package (39). Venn diagrams were constructed using Venny (version 2.1.0) online freeware and statistical comparisons were performed with GraphPad Prism (version 6).

#### Biofilm formation

Small pieces of silicon (approx. 1.5cm by 1.5cm size) were cut from silicone tubing, washed with water, dried and autoclaved. The silicone pieces were incubated for overnight with Fetal Bovine Serum (Gibco) and then washed with PBS (Phosphate Buffered Saline).

A loop of the wild type strain (SC5314) and *bta1*ΔΔ mutant were washed twice with PBS and then diluted to OD_600_-0.5 in 2ml of spider, Lee’s, RPMI1640 and DMEM media in a tissue culture treated 24 well plate (Corning). The silicone pieces were then added to the media and kept for incubation for 90min at 37°C and 150 rpm. After the initial attachment of the yeast cells, the pieces were gently washed with PBS to remove the non-adherent cells. Fresh media were added to each well and incubated for overnight at 37°C with 150rpm. Biofilms were further observed by confocal laser scanning microscopy.

#### Confocal Laser Scanning Microscopy (CLSM)

Biofilms were washed twice with PBS and live stained with 1mg/ml calcofluor white (which stains fungal cell wall) for 30 min at 37°C. Cells were again washed gently with PBS to remove excess stain and placed onto glass microscope slide. Calcofluor-white is normally imaged with a single excitation and emission, but using two channels provided better contrast in later visualizations. The excitation laser lines of 405 nm and 488 nm were used and the emissions were 425-475 nm and 500-550 nm respectively.

Confocal laser scanning microscopy was performed with a Nikon A1 CLSM mounted on a Nikon 90i compound microscope and controlled by NIS-Elements (4.40). Samples were imaged in PBS at room temperature with a Nikon CFIPl an Apochromat 60x water immersion objective (NA 1.20). The biofilms had characteristic morphologies, thus data (Biofilm thickness) was collected in the z-axis at 1 um steps from top to bottom. The lower limits were based on signal drop off due to dense samples’ refraction or samples, which were thicker than the total working distance of the lens. After acquisition, z-series data was pseudocolored (depth coded) and projected as a 3D image with NIS-Elements.

#### Amino Acid Profiling by Ultra-Performance Liquid Chromatography (UPLC)

The wild type SN152 was grown aerobically at 30°C to A_600_ 0.8-1.0 in phosphate free synthetic complete media containing high (2 mM) or low (50 µM) phosphate concentrations and frozen in liquid nitrogen. The samples for free amino acid (FAA) analysis were processed as described by Hacham *et al.* (40). Briefly, frozen samples (10ml each) were mechanically lysed by glass beads (acid washed) in the presence of 600 μL of water:chloroform:methanol (3:5:12, v/v). Centrifuge and supernatant was collected followed by re-extraction of the residue with another 600 μL of water:chloroform:methanol (3:5:12, v/v). The supernatants were pooled. Chloroform (300 μL) and water (450 μL) were added, and the resulting mixture was centrifuged again. The upper water:methanol layer was collected and dried. Finally the samples were resuspended in HCl 20 mM and further used for total free amino acid quantification on an Agilent 1290 Infinity II UPLC.

The extracted amino acids were derivatized using AccQ-Tag reagents (Waters, Milford, MA, USA) according to the manufacturer’s protocol. Briefly, 70 μL of AccQ-Tag Ultra borate buffer was added to 10 μL of the biological extract to ensure optimum pH. 20 μL of AccQ-Tag reagent previously dissolved in 1.0 mL of AccQ-Tag Ultra reagent diluent were added to each sample and the tubes were incubated at 55 °C for 10 min. The same steps were followed for the external standard calibration curve. Liquid chromatographic separation was performed on the AccQ-Tag Ultra column (Waters, 2.1 mm I.D. × 100 mm, 1.7 μm) at 43°C with a 1290 Infinity II UPLC system (Agilent, Santa Clara, CA, USA). The UPLC is equipped with the Flexible quaternary pump running at a flow rate of 0.7 mL/min. Mobile phases A-D are: A=100% Waters AccQ-Tag Eluent A; B=10% Waters AccQ-Tag Eluent B; C=100% Milli-Q water; D=100% Waters AccQ-Tag Eluent B. The gradient was as follows : T=0, 9.9% A, 90.1% C; T=0.29 min, 9.9% A, 90.1% C; T=4.84 min, 9.1% A, 70% B, 20.9% C, 0% D; T=6.45 min, 8% A, 15.6% B, 58.9% C, 17.5% D; T=6.65 min, 8% A, 15.6% B, 58.9% C, 17.5% D; T=7.04 min, 7.8% A, 0% B, 71.9% C, 20.3% D; T=7.64 min, 13.7% A, 0% B, 36.3% C, 50% D; T=8.89 min, 13.7% A, 0% B, 36.3% C, 50% D; T=8.98 min, 9.9% A, 0% B, 90.1% C, 0% D. The signal was detected using the PDA detector at 260 nm with a sampling rate of 40 Hz. One microliter of sample and standards were injected for analysis. The concentrations of amino acids were calculated using the external standard calibration curve. The concentrations in pmol/μL for each amino acid were detected in each sample and are calculated using a series of standard dilutions run before the samples.

**Fig. 9.**
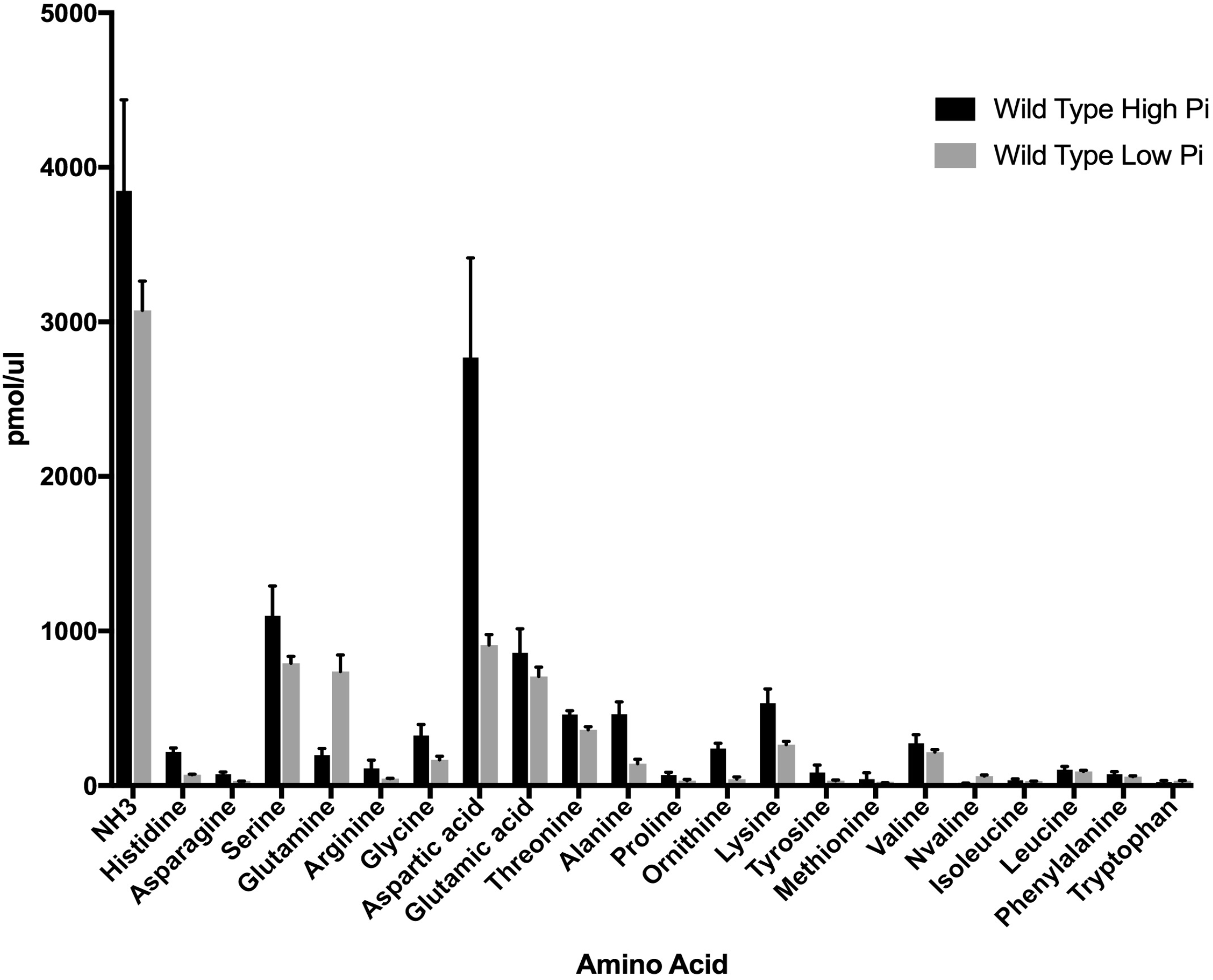
Amino acid analysis of free amino acids in P_*i*_ replete and limited cells. Cultures grown with ample (2 mM) or limiting (50 μM) initial P_*i*_ concentrations were lysed and aqueous extracts were prepared as described in Materials and Methods, and the amino acids quantified by UPLC.

